# TCMCard: A High-Confidence Digital Infrastructure for Traditional Chinese Medicine Quantified by Multi-Dimensional Evidence Integration

**DOI:** 10.64898/2026.04.07.716940

**Authors:** Yuqi Wang, Wenqing Dong, Junkai Yao, Kai Wang, Leyuan Zhang, Yuanrong Wang, Shanshan Guo, Haorui Li, Hengjia Cai, Xiting Wang, Yu Li

## Abstract

Network pharmacology has become a widely used approach for deciphering multi-component, multi-target mechanisms of traditional Chinese medicine (TCM). Here we introduce TCMCard, a high-confidence digital infrastructure built on a Multi-Dimensional Evidence Integration (MDEI) framework. The framework integrates experimental activity data from authoritative chemical databases, literature-derived evidence, and structure-based similarity inference. Preprocessing steps include chemical structure normalization, species-specific filtering, and target quality scoring. Applied to conventional interaction datasets, this pipeline leads to the removal of over 60% of low-confidence noise. TCMCard supports network pharmacology exploration through an interactive visualization platform, and module analysis identifies functionally relevant communities that offer insights into the synergistic actions of TCM formulas. Overall, TCMCard may help move the field beyond simple data aggregation toward evidence-informed curation and quality-driven analysis. As an interactive and publicly accessible platform, it reveals an organized backbone within complex interaction networks, offering a more reliable basis for understanding multi-component synergy in TCM.

## Introduction

Traditional Chinese Medicine (TCM) is a highly intricate therapeutic system characterized by multi-component interventions, multi-target modulation, and context-dependent clinical effects. Instead of acting through single dominant targets, TCM formulas are believed to provide therapeutic benefits by simultaneously perturbing interconnected molecular processes across multiple biological scales[1]. This systems-level approach closely aligns with modern network biology, in which disease and treatment are increasingly viewed as emergent properties of molecular interaction networks rather than as isolated linear pathways. Consequently, over the past two decades, network pharmacology has become the leading framework for understanding the molecular mechanisms underlying TCM formulas[2, 3]. The rapid growth of this field relies heavily on the parallel development of TCM-focused bioinformatics resources, which now serve as the core data infrastructure for large-scale computational analysis and mechanism inference[4].

Over time, TCM databases have evolved from simple repositories of herbal knowledge into increasingly diverse platforms that support chemical annotation, target prediction, symptom mapping, and network-based analysis[5]. Early resources such as TCM-ID and TCM Database@Taiwan primarily focused on digitizing medicinal materials and phytochemical structures[6, 7], laying the foundation for virtual screening and compound-focused research. The subsequent rise of systems pharmacology shifted database development toward integrated herb–ingredient-target–disease frameworks, represented by platforms such as TCMSP[8] and TCMID[9]. This analytical paradigm was further extended by BATMAN-TCM[10] and HIT 2.0[11], which strengthened the ingredient – target evidence layer through prediction and literature curation, respectively. In parallel, TCM-Mesh[12] and YaTCM[13] integrated heterogeneous datasets with built-in tools for network pharmacology and pathway analysis. More comprehensive databases then appeared, represented by ETCM[14], SymMap[15], HERB[16], TCMBank[17], and SuperTCM[18], further expanding this landscape through symptom-level translation, experimental and clinical evidence integration, large-scale data aggregation, and cross-resource identifier harmonization. Collectively, these resources have greatly advanced TCM systems pharmacology by enabling more integrative and interconnected analyses across formulas, herbs, ingredients, targets, phenotypes, and layers of molecular evidence.

However, the growth of TCM informatics has been driven more by increasing data volume than by systematic efforts to improve evidence quality[19]. Many modern platforms inherit, merge, or repurpose associations from earlier databases such as TCMSP, where foundational ingredient–target relationships often still originate from text mining, structural similarity inference, or outdated prediction pipelines developed before modern bioactivity resources became widely available[20, 21]. Consequently, expanding the database does not always improve biological reliability. Instead, low-confidence or weakly validated associations are often repeatedly circulated across platforms, creating dense, loosely constrained interaction networks in which distinguishing signal from noise is challenging[21, 22]. This issue is especially critical in TCM network pharmacology, where downstream analyses such as enrichment testing, hub prioritization, and pathway interpretation are highly affected by redundancy, variable confidence level, and false-positive inflation[20, 23]. As TCM databases evolve toward more mechanism-based and experimentally verifiable models, evidence quality must be reassessed along with network expansion. Otherwise, larger networks could hinder interpretability rather than enhance it. Simultaneously, transparency and reproducibility of evidence remain significant unmet needs[24].

These limitations call for a shift from coverage-focused aggregation to evidence-based curation and confidence-based stratification. Removing unreliable relationships selectively may be more effective than randomly expanding the network topology. Here, we developed TCM Comprehensive Analysis and Reliable Database (TCMCard), a curated and analytically manageable TCM knowledge framework based on a formula-centered and evidence-weighted approach. Instead of starting with the broadest inclusion of all available entities, we initially defined a high-quality core formula–herb space rooted in authoritative sources, including the Pharmacopoeia of the People’s Republic of China (2020 edition) (PC 2020) and well-known ancient prescriptions[25]. We then refined the ingredient layer through PC 2020 cross-referencing and cheminformatic quality checks, systematically reevaluating and expanding the ingredient-target relationships using modern high-confidence sources from ChEMBL and PubChem BioAssay[26, 27].

To assess the quality of evidence across the resulting network, especially for ingredient–target interactions, we developed a Multi-Dimensional Evidence Integration (MDEI) framework and eliminated over 60% of low-confidence connections from traditional datasets. Based on this, we built a quality-stratified TCM knowledge graph that links formulas, herbs, ingredients, targets, and disease phenotypes. Using this curated network, we further explored systems-level properties of TCM, such as pharmacological hub structure, pleiotropic convergence, and the scaling relationships between formula size and molecular complexity. Finally, we integrated these data resources and analytical workflows into TCMCard (https://www.tcmcard.com), an interactive web platform for standardized network pharmacology, knowledge retrieval, and predictive inference. Overall, this work offers both a curated resource and a quantitative framework to advance mechanism-focused TCM research based on higher-confidence evidence.

## Materials and methods

### 2.1 Construction of a Curated Core Formula and Herb Dataset

To construct a pharmacologically interpretable and topologically coherent TCM knowledge graph (TCM-KG), we applied a cascading filtering strategy to define a curated core dataset. The dataset construction followed a formula-centric design, in which herbs were retained only if they occurred in validated formulas, thereby ensuring a biologically contextualized and internally connected network.

Formulas composed of a single herb were excluded to preserve combinatorial structures suitable for compatibility and synergy analysis. Redundant dosage-form records for the same prescription, such as capsules and tablets, were consolidated into a single representative entry. Only formulas explicitly documented in authoritative sources, including the PC 2020 or recognized ancient classical prescriptions, were retained. Proprietary preparations without standardized clinical lineage were excluded. In addition, formulas for which more than 50% of constituent herbs lacked verifiable chemical or target annotations were removed to improve downstream interpretability and reduce low-confidence noise. Synonymous formula names were further merged through deep deduplication.

After establishing the core formula set, herbs were added through a series of inclusion criteria. Only unique herbs found in the validated core formulas were kept. This approach excluded isolated herbal entities and made sure that all herb connections were based on a confirmed therapeutic context.

### 2.2 Validation and Refinement of Chemical Ingredients

To ensure the chemical validity and biological relevance of compounds, we implemented a multi-dimensional verification strategy that integrated internal database mining, external pharmacological cross-referencing, and cheminformatic filtering for all herbs retained in the core herb dataset.

All small-molecule ingredients explicitly linked to the core herb set were initially compiled from the primary repository, ensuring each ingredient had a traceable botanical origin. To assess chemical reliability, we built an Artificial Intelligence (AI) agentic pipeline to extract quality-control markers and index components from the digital edition of PC2020. These pharmacopoeial components were then cross-checked against the combined ingredient collection to label high-confidence marker compounds and differentiate them from generic or trace constituents.

To reduce chemical noise and experimental artifacts, a stringent RDKit-based cheminformatic pipeline was applied[28]. All SMILES strings were validated to remove entries with invalid valence states or incomplete structural representations. Pan-Assay Interference Compounds (PAINS) filtering was performed to identify and flag potential assay-interfering structures[29]. Key physicochemical properties, including molecular weight, LogP, and a quantitative estimate of drug-likeness (QED), were further calculated to characterize drug-like features[30]. Finally, ingredient distribution across the herbal network was examined to identify and exclude ubiquitous, non-specific metabolites, such as sucrose, starch, and common fatty acids that were unlikely to contribute specific pharmacological signals (Fig. 1A).

**Fig. 1.**
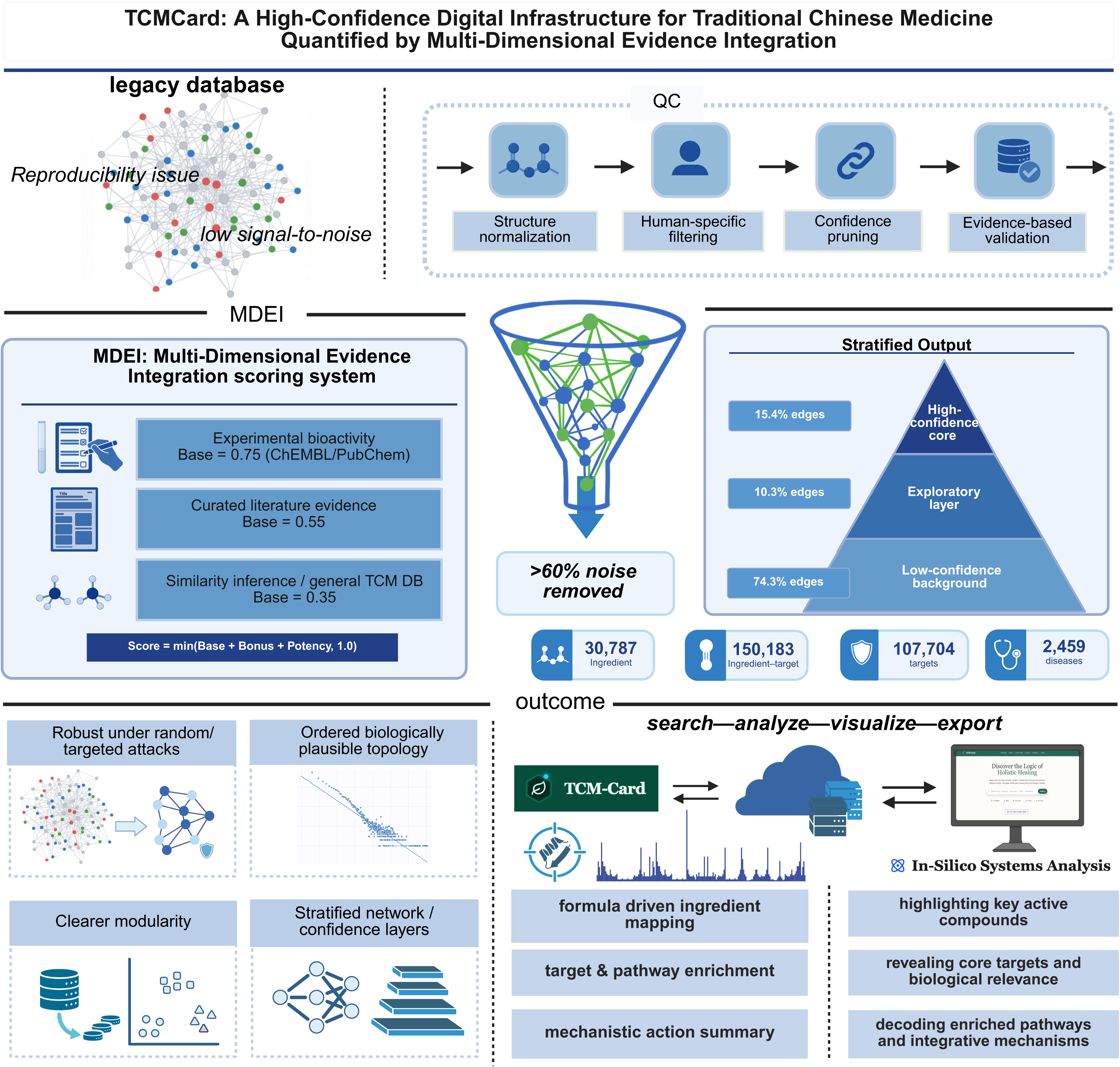
Systematic architecture of TCMCard for data integration, evidence-aware curation, and construction of the high-confidence TCM-KG. The TCMCard framework transforms heterogeneous Traditional Chinese Medicine (TCM) data into an evidence-weighted knowledge graph through sequential integration, quality control, and network assembly. (A) Multi-source data acquisition. Integration of heterogeneous inputs from TCM resources, chemical bioassay databases (ChEMBL, PubChem), and clinical repositories. (B) Standardized curation and preprocessing. Implementation of species-specific restriction (Homo sapiens), structure normalization, and the Synergistic Quality Scoring (SQS) algorithm to establish a high-fidelity core. (C) Multi-Dimensional Evidence Integration (MDEI). Interaction reliability is quantified by an automated scoring engine that synthesizes evidence provenance, cross-source consensus, and pChEMBL-derived potency bonuses. (D) Multi-level biological validation strategy. Systematic assessment of entity quality, relationship confidence, and network robustness via the Multi-dimensional Completeness Index (MCI) and topological stability analysis.(E) Resulting in a high-resolution TCM-KG. The final curated knowledge graph organized into formula–herb– ingredient, functional interactome, and clinical phenotype layers, featuring evidence-weighted connectivity and validated multi-layer architecture.

### 2.3. Optimization and Validation of Ingredient-Target Interactions

To construct a robust ingredient-target network, we prioritized reliability over quantity and implemented a dual strategy of validation and expansion using ChEMBL (v33) and PubChem BioAssay. Consolidated interaction data were standardized by retaining only Homo sapiens targets and categorizing evidence as database-derived or literature-supported. Core ingredients were mapped to both databases, and interactions with pChEMBL ≥ 5 or equivalent activity values <10 μM were classified as experimentally validated (Fig. 1B). These resources were further used to identify high-affinity interactions absent from conventional TCM databases, which were incorporated as activity-based extensions.

### 2.4 Systematic Integration of Disease Phenotypes

Clinical relevance served as the main criterion for collecting and filtering disease-related data, ensuring that the final interactome reflected validated disease-molecular connections. Disease association data were collected at scale using a custom scraping bot to retrieve more than 30,000 disease records from the HERB database[31]. The automated pipeline extracted and harmonized diverse relationships, including disease-herb, disease-formula, disease-ingredient, and disease-target associations, into standardized analytical formats. Using this clinical evidence layer, a reverse-validation approach was applied to select the disease node. Disease nodes were retained only if they showed statistically significant associations, after Benjamini–Hochberg false discovery rate (FDR BH) correction. They also had to be linked to at least one verified entity in the curated core dataset, such as validated formulas, core herbs, refined ingredients, or prioritized targets. This procedure ensured that each disease node in the knowledge graph was mechanistically linked to a verified TCM molecular framework rather than being included as an isolated association.

### 2.5 Multi-Dimensional Evidence Integration for Relationship Quality Assessment

To support detailed mechanistic analyses, we developed an MDEI framework to assess the confidence in ingredient-target interactions. This framework combined three separate aspects of evidence quality: source hierarchy, cross-platform reproducibility, and quantitative bioactivity strength.

First, a hierarchical baseline score was assigned based on data provenance. Interactions derived from quantitative bioassays in ChEMBL or PubChem were given a baseline score of 0.75. Interactions from expert-curated literature received a score of 0.55, while entries from general TCM databases were conservatively weighted at 0.35. Second, to account for reproducibility across different resources, a multi-source consensus bonus was added: interactions supported by two or three independent sources gained an extra +0.10 or +0.15, respectively. Third, for interactions with quantitative potency data such as pChEMBL values, a continuous potency bonus was used to prioritize higher-affinity binding events: potency bonuses = min (0.05 × (pChEMBL − 5), 0.20). This approach distinguishes nanomolar-range binders (typically pChEMBL ≥ 8, corresponding to sub-10 nM activity) from micromolar-range associations. The final confidence score was calculated as the sum of the baseline score, consensus bonus, and potency bonus, capped at 1.0 (Fig. 1C). This design maintains a broad dynamic range while allowing for strict prioritization of high-confidence evidence.

### 2.6 Comprehensive Data Quality Assessment Framework

To assess the reliability of the underlying digital infrastructure, we established a Multi-dimensional Completeness Index (MCI) across five core entity types: Formula, Herb, Ingredient, Target, and Disease. The MCI integrated three dimensions of data quality: information completeness, cross-database validation, and structural integrity. Information completeness was evaluated by the fill rate of essential attributes, such as nomenclature and identifiers. Cross-database validation quantified the degree of corroboration among authoritative sources. Structural integrity, particularly for chemical ingredients, included SMILES validation and standardization. Entities with composite scores ≥0.80 were classified as high quality, whereas those scoring between 0.50 and 0.80 were classified as medium quality.

For relationships other than ingredient-target associations, we implemented an evidence consensus model. For compositional relationships, confidence was anchored to authoritative sources such as the PC2020 and specialized TCM repositories, and then weighted according to agreement across independent datasets. For clinical and bio-effect relationships, the model distinguished experimentally or clinically validated associations from text-mined records and assigned higher confidence to relationships supported by clinical reports or jointly represented in multiple biological databases (Fig. 1D).

### 2.7 Quantitative Characterization of Global Network Topology and Biological Scaling Features

To quantify the systemic properties of TCM compatibility, we analyzed the TCM-KG using network topology metrics and complexity scaling relationships. For hub-target synergy analysis, degree centrality was calculated for all components to identify central pharmacological hubs mediating convergent multi-ingredient effects. Interaction strengths across network layers were weighted using the MDEI framework to construct an evidence-informed interaction landscape. To evaluate formula complexity and pharmacological scaling, nonlinear regression was performed to relate botanical diversity (number of herbs) to molecular diversity (number of unique ingredients and targets) at the formula level. A power-law model was fitted to characterize how molecular complexity scaled with prescription size. In addition, probability density functions and kernel density estimation were used to analyze the distributions of formula skeletons, aiming to describe topological patterns underlying the transition from simple combinations to complex multi-component systems.

To reduce dimensionality and noise in the multi-component network, we developed a synergistic quality scoring (SQS) algorithm to prioritize core targets. The score for a target twas defined as

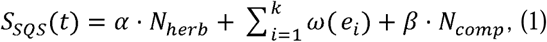

where N_herb_ denotes the number of distinct herbs converging on the target, w(e_i_) denotes the confidence weight of each ingredient-target interaction, and N_comp_ denotes the number of active ingredients linked to the target. A dominant weight was assigned to N_herb_(a = 10.0) and a minor auxiliary weight to N_comp_(f3 = 0.1). For each interaction edge w(e_i_) =1.5 were assigned when the evidence score exceeded 0.8, indicating experimental validation or high-confidence prediction; otherwise, edge w(e_i_) = 0.5. Targets were ranked by SQS, and the top 50 were retained for downstream analyses.

### 2.8 Platform Architecture and Implementation

TCMCard is a lightweight web platform designed to improve accessibility for researchers without advanced computational expertise. The platform was implemented as a responsive single-page application, with a frontend developed using React.js and Vite and a backend built with Python (FastAPI). Computationally intensive tasks, including pathway enrichment analyses, were executed using integrated Python libraries such as GSEApy and Pandas[32]. A central component of the platform is an interactive network visualization engine built on D3.js, which supports real-time exploration through functions such as zooming, node dragging, and hierarchical rendering of multi-layer networks spanning herbs, ingredients, targets, and diseases. The entire system was containerized to ensure cross-platform compatibility and facilitate deployment on both cloud servers and local workstations.

## Results

### 3.1 Construction and Validation of a Curated TCM Target-Disease Landscape

We constructed a curated TCM landscape by refining over 6,700 entries into a high-quality core set of 1,774 standardized formulas. While TCMCard retains the multi-entity integration common to existing TCM resources, it is distinguished by formula-centered curation, evidence-prioritized refinement, and confidence-stratified relationship assessment (Table S1). This process focused on canonical sources, including PC2020 and recognized ancient classical prescriptions, while removing 478 single-herb isolates and extensive commercial redundancies. The resulting dataset was reduced to 748 unique herbs and 30,787 chemical ingredients linked via 56,678 herb-ingredient relationships. Cross-referencing with PC2020 confirmed that this chemical space maintained key therapeutic components and exhibited high structural consistency. A clinical phenotype layer was subsequently incorporated, comprising 2,459 unique disease nodes connected through 133,711 validated disease-associated edges.

Mapping this curated chemical layer to the functional interactome identified 150,183 ingredient-target interactions involving 10,704 human targets. Of these, 15,874 interactions (10.6%) were directly supported by high-activity experimental evidence from ChEMBL and PubChem (Fig. 1E). Additionally, targeted mining of core ingredients identified 7,181 high-affinity interactions not found in traditional TCM databases but validated by modern bioactivity assays, thus expanding the legacy knowledge base with experimentally supported associations (Table S2). Collectively, these layers integrated formulas, herbs, ingredients, targets, and disease information into a unified knowledge framework and established a multi-layer TCM target–disease landscape, laying the groundwork for subsequent confidence stratification, topological analysis, and platform-level implementation (Fig. 2A-B).

**Fig. 2.**
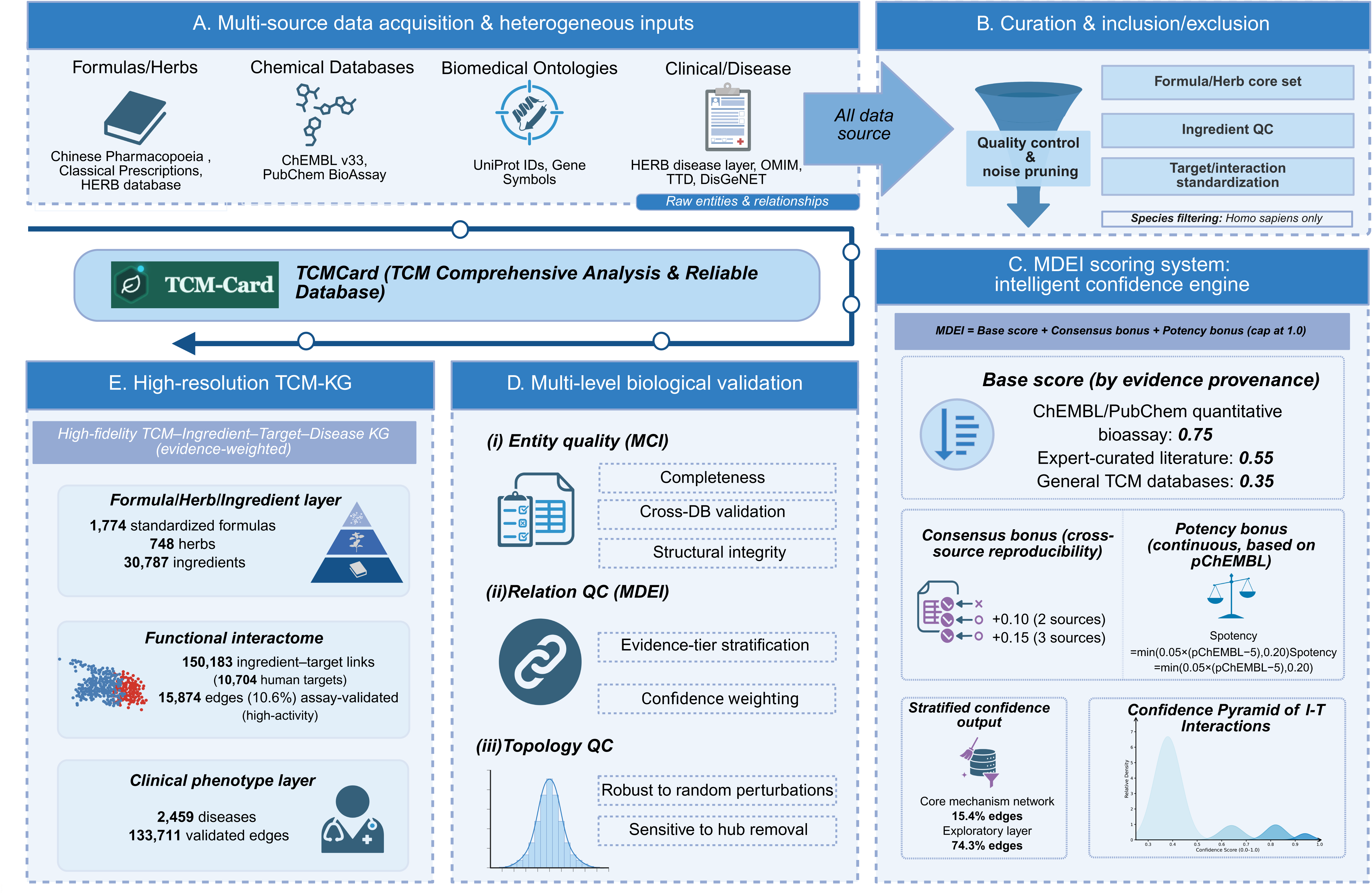
Global composition, confidence stratification, and topological scaling laws of the TCM-KG. Comprehensive quantitative analysis of the structural integrity and organizational logic governing the knowledge graph. (A) TCM-KG global landscape with multi-layered topology and distribution of major entity types. (B) Quantitative breakdown of nodes and interactions across the integrated categories in the network composition. (C) Scale-free architecture following a power-law fit (y = 1.24), characteristic of hub-dominated networks. (D) Key target nodes weighted by the number of ingredient interactions and MDEI confidence score. (E) Top multi-target molecules ranked by biological coverage. (F) Frequency distribution of herb counts per prescription (median = 8). (G) Scaling relationship between botanical size and molecular diversity, revealing an asymptotic saturation (R^2^ = 0.79). (H) Distribution of ingredient diversity across formula size categories exhibits a Chemical complexity scaling.

### 3.2 Multidimensional Quality Assessment and Structural Integrity of the TCM-KG

To assess the integrity of the TCM-KG, we implemented a multi-step curation and standardized workflow that converts diverse raw interactions into a unified knowledge base. After species-specific filtering and identifier normalization, the MDEI scoring system was applied to categorize interaction quality across various evidence levels. The score distribution revealed four distinct peaks instead of a smooth range. This pattern clearly differentiated evidence-supported, high-confidence interactions from lower-confidence background associations. At the same time, it maintained a concise core of consolidated TCM knowledge and prioritized pharmacological inference.

Systematic assessment of the 46,472 entity nodes showed high overall data completeness and consistency, with 99.8% of entities classified as high quality (Table S3). This fidelity was supported by category-specific quality control measures, including SMILES-based standardization and structural verification for all 30,787 ingredients, 100% canonical annotation coverage for the 10,704 human targets, and verified taxonomic origins with bilingual nomenclature for all 748 herb nodes.

Application of the MDEI framework revealed a hierarchical confidence distribution across the integrated network. A minority of ingredient-target interactions, accounting for 15.4% of all edges, formed a high-confidence core mechanism network (Table S4), dominated by experimentally validated targets with nanomolar potency. The majority of the landscape, accounting for 74.3% of edges, remained in the lower-confidence tier, representing the exploratory search space. This distribution is consistent with the stringent scoring criteria of the MDEI model and indicates that the framework effectively distinguishes a compact evidence-supported core from a much larger set of heterogeneous but potentially informative associations.

Confidence scoring of the unified edges revealed a clear hierarchical structure (Table 1). Compositional relationships showed the highest mean scores, including herb–ingredient (0.750) and formula–herb (0.700) edges, indicating robust mapping from prescriptions to botanical components and curated phytochemical constituents. Herb–disease, formula–disease, and target–disease relationships formed an intermediate tier (all 0.600), whereas ingredient–disease (0.500) and ingredient–target (0.460) edges occupied a broader, lower-confidence layer.

**Table 1.**
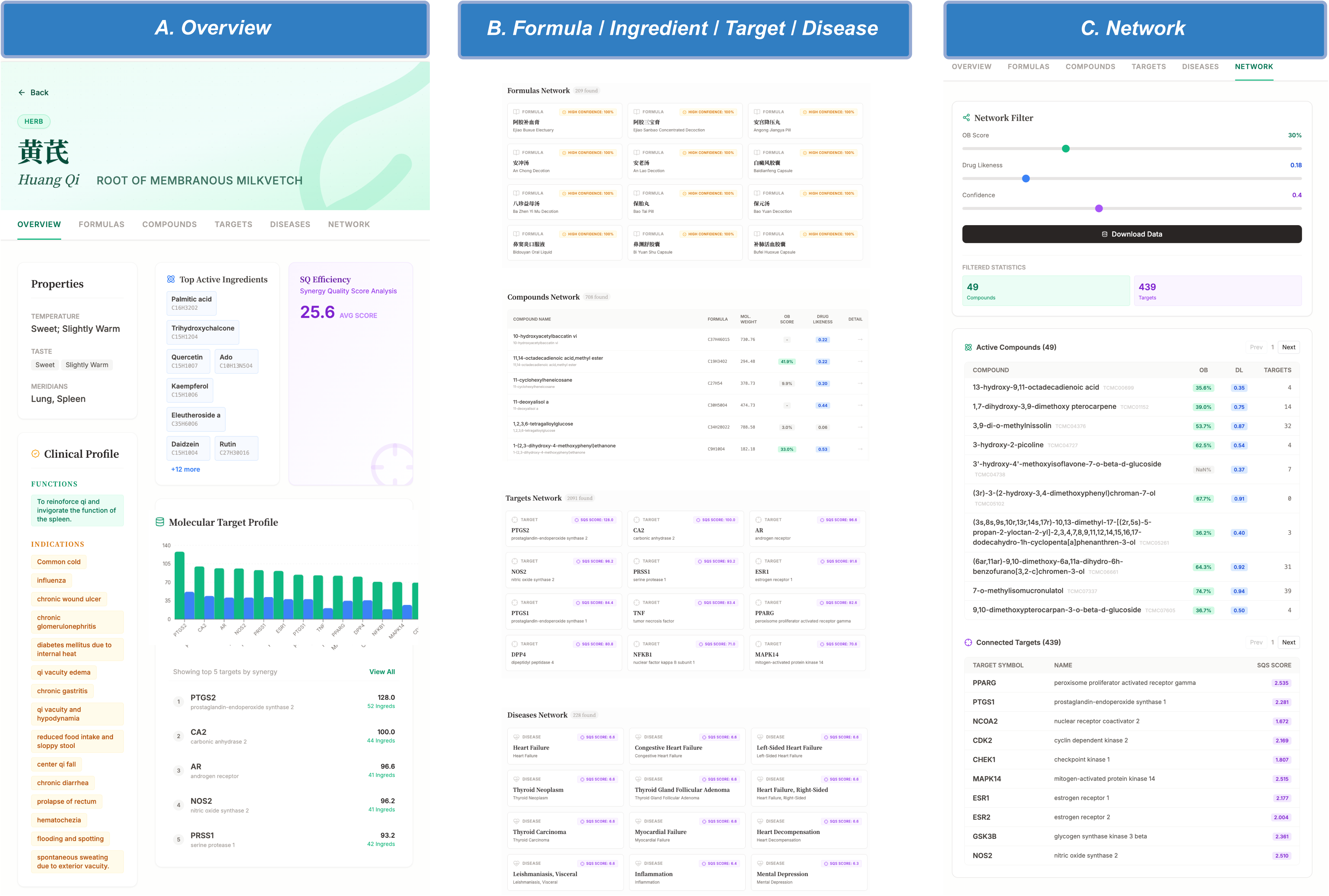
Hierarchical quality distribution and semantic value of relational tiers.

Given their large edge counts, these lower-scoring tiers likely represent a more exploratory but still structured search space for downstream analysis. Together, these results indicate that the TCM-KG provides a structurally coherent and quantitatively stratified framework for network-based interrogation of TCM mechanisms.

### 3.3 Hierarchical Nodal Orchestration and Pleiotropic Synergy of Pharmacological Hubs

Analysis of the TCM-KG revealed a hierarchical organization of pharmacological hubs, wherein upstream hubs orchestrate downstream effectors and exhibit pleiotropic synergy that collectively contributes to formula efficacy. The network exhibited a scale-free architecture with power-law degree distribution (Fig. 2C), indicating that connectivity is dominated by a few highly connected nodes. Robustness analysis confirmed that the network tolerates random perturbations but is vulnerable to targeted disruption of high-degree nodes, underscoring the critical role of hubs in maintaining network integrity.

By integrating degree centrality with MDEI-weighted interaction profiles, we identified targets such as PTGS2 and AKR1B1 as major convergence points in the pharmacological landscape. These targets linked diverse ingredients to shared pathological processes across multiple disease contexts (Fig. 2D). At the molecular layer, pleiotropic ingredients including quercetin and sitosterol emerged as broad-spectrum regulators connected to more than 200 targets (Fig. 2E). Notably, several of these central hubs were enriched within the higher-confidence core mechanism layer defined by MDEI stratification, linking network centrality to stronger evidence support.

Together, these observations suggest that the TCM-KG is organized around hierarchically distributed hubs spanning both target and ingredient layers. This architecture provides a systems-level explanation for how multi-component TCM interventions may achieve coordinated effects by converging on a limited set of biologically influential nodes rather than through diffuse, uniformly distributed associations.

### 3.4 Scaling Relationships Governing Formula Complexity in the TCM-KG

Beyond identifying individual hubs, we next examined whether formula organization within the TCM-KG adhered to broader quantitative scaling principles. Analysis of formula composition showed that the median number of herbs per prescription was 8 (Fig. 2F), indicating that moderate combinatorial complexity predominates across the curated formula space. We then assessed how increases in botanical diversity related to changes in ingredient-level complexity. Power-law regression revealed a non-linear scaling relationship in which the expansion of ingredient and target space followed a pattern of diminishing returns (Fig. 2G). In particular, the scaling trajectory approached an asymptotic regime at approximately 15 herbs (Fig. 2H), beyond which additional herbs contributed relatively little to unique ingredient coverage and instead increased functional redundancy.

These findings suggest that formula composition in the curated TCM-KG is not random but is constrained by an efficiency-like scaling rule that balances breadth of coverage against increasing redundancy. This quantitative pattern provides a framework for interpreting TCM prescription complexity as a trade-off between synergistic coverage and molecular efficiency.

### 3.5 TCMCard: a cloud-based analytical platform for scalable and reproducible knowledge discovery

To translate the curated TCM-KG into an accessible research resource, we developed TCMCard (https://www.tcmcard.com), a web-based platform that integrates knowledge retrieval, network analysis, and predictive inference within a unified analytical environment (Fig. 3A). Unlike conventional repositories focused on static record browsing, TCMCard functions as a dynamic analytical engine that supports reproducible, workflow-oriented interrogation of TCM molecular mechanisms at scale.

**Fig. 3.**
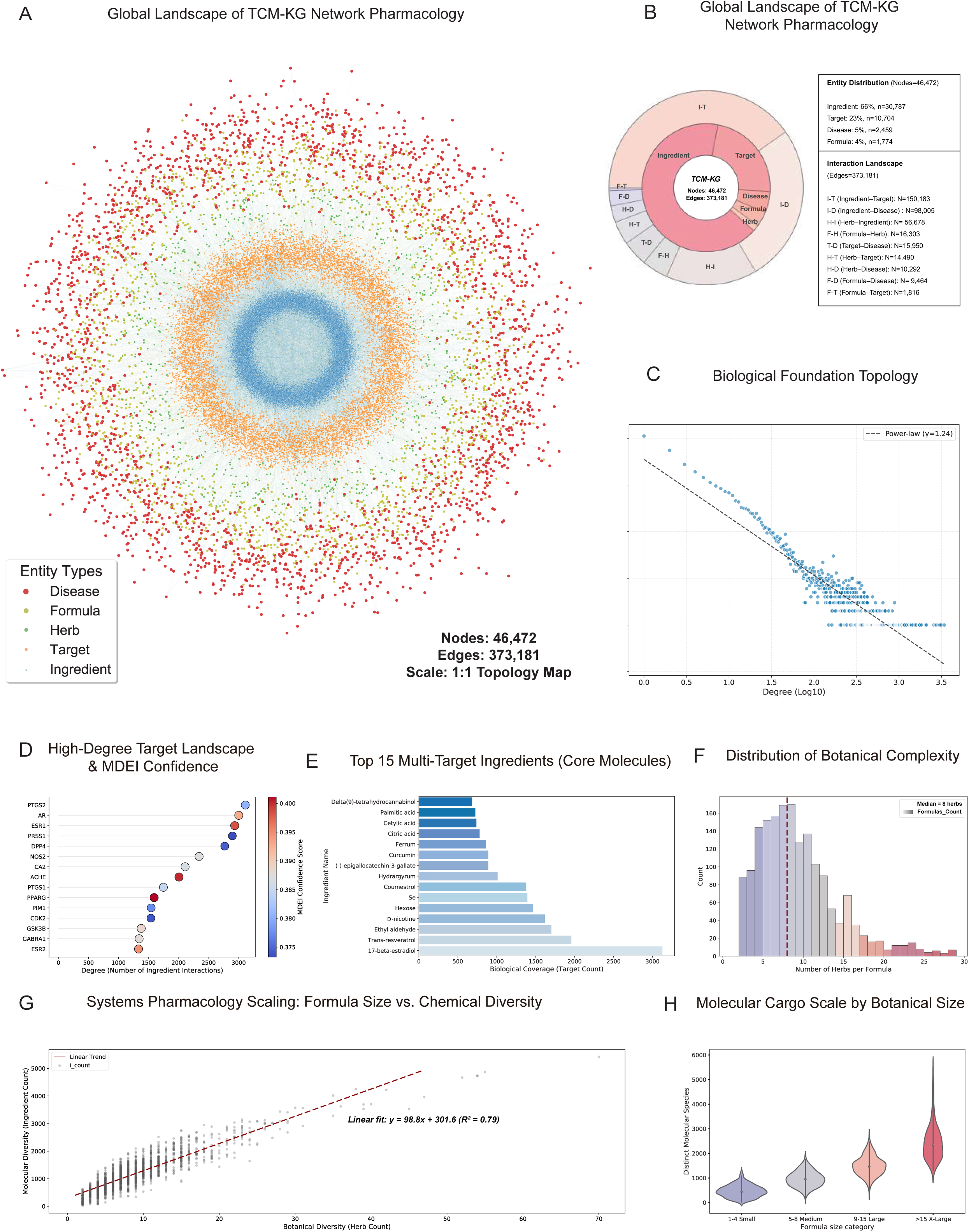
Cloud-native infrastructure and AI-driven predictive analytics of TCMCard. The platform is engineered as a scalable ecosystem that leverages cloud-native computing and deep learning to facilitate knowledge discovery. (A) Cloud-native ecosystem featuring a React-based interactive frontend and a dual-backend framework for session management and intensive computation. The infrastructure supports sub-second query latency across 373,181 edges. (B) AI-powered inference framework utilizing the MDEI-weighted interactome as a high-quality substrate. The engine supports deep-learning-based target prediction, ingredient affinity scoring, and drug repositioning analytics to prioritize candidates for experimental validation.

TCMCard is built on a scalable web architecture that enables high-throughput, low-latency access to the TCM-KG, which contains 373,181 edges. This infrastructure supports efficient exploration of large-scale graph relationships and provides a streamlined interface for querying formulas, herbs, ingredients, targets, and diseases.

Beyond descriptive analysis, TCMCard incorporates AI-powered modules to support predictive exploration of uncharacterized pharmacological relationships. Using the MDEI-weighted interactome as a structured, evidence-informed substrate, the platform enables target prediction, ingredient–target affinity scoring, and candidate indication expansion (Fig. 3B). These predictive functions allow users to move from retrospective knowledge retrieval to forward-looking hypothesis generation. By integrating curated graph knowledge with machine learning-based inference, TCMCard provides a practical framework for prioritizing candidate mechanisms and guiding experimental follow-up in TCM modernization research.

### 3.6 Case Study: Dissecting Formula Mechanisms Using TCMCard

To demonstrate the practical application of TCMCard in mechanism-oriented analysis, we selected Liuwei Dihuang Pill (LDP) as an example of formula-centered analysis and Astragali radix as an example of entity-centered knowledge navigation. These case studies illustrate how users can move from entry-level retrieval to downstream mechanistic interpretation through the integrated analytical modules of the platform.

The LDP case illustrates the web-based workflow from formula retrieval to downstream mechanistic interpretation. Users can access the corresponding formula page by selecting the formula entry or by searching for “Liuwei Dihuang Pill” (Fig. 4A). The interface presents structured information for the selected prescription, including category information, formulation composition, clinical profile, and system pharmacology-related modules. In addition to overview-level content, the page also provides dedicated subpages for herbs, compounds, targets, diseases, and network exploration, allowing users to examine the multi-layer organization of a formula within the TCM-KG.

**Fig. 4.**
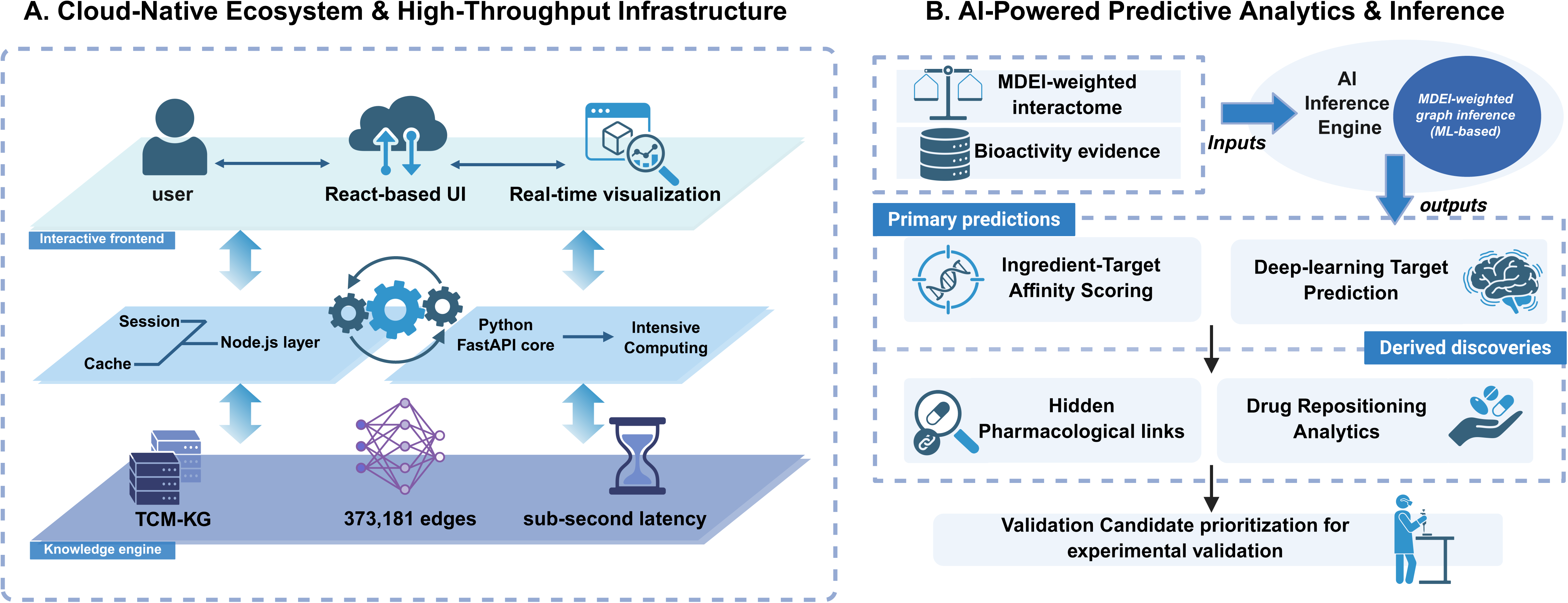
Automated analytical workflows and multi-entity interactive interface modules. User-centric environment integrating standardized computational protocols with multi-dimensional data exploration. (A) Standardized network pharmacology pipeline. End-to-end workflow enabling automated ingredient mapping, target enrichment, and pathway crosstalk analysis via a single-click interface to ensure reproducibility. (B) Integrated knowledge exploration interfaces. Database modules providing detail pages for herbs, compounds, targets, and diseases, incorporating entity profiles, MDEI-weighted relationships, and AI-driven mechanism interpretations.

To further illustrate formula analysis in a disease-specific context, LDP and diabetes were selected as an example pair. The resulting workflow includes ingredient mapping and network construction (Fig. 4B), target-related summary and enrichment analysis (Fig. 4C), and pathway enrichment analysis with selectable reference libraries (Fig. 4D). Together, these modules provide a stepwise view of formula-associated components, inferred targets, and enriched biological pathways, thereby supporting investigation of potential pharmacological mechanisms. To improve interpretability, TCMCard also provides an AI-assisted reporting module that automatically summarizes the analytical results into a readable mechanism report (Fig. 4E). The generated report integrates multiple layers of evidence into sections such as executive summary, key active compounds, molecular mechanisms, core target analysis, enriched pathway analysis, and conclusion, enabling users to move from raw analytical outputs to clearer mechanistic hypotheses.

At the entity level, TCMCard also provides dedicated pages for herbs, compounds, targets, and diseases. Figure 5 presents Astragali radix as a representative example of herb-centered exploration. The page includes three main components: an overview section, entity-specific relationship modules, and a network exploration page. The overview section summarizes key information for the selected herb, including basic properties, clinical profile, top active ingredients, synergy quality score, and molecular target profile (Fig. 5A). Related formulas, compounds, targets, and diseases are displayed through entity-specific modules (Fig. 5B). The network page further supports filtered exploration of herb-associated active compounds and connected targets using thresholds for oral bioavailability, drug-likeness, and confidence, with downloadable results available for further analysis (Fig. 5C). Together, these modules demonstrate how TCMCard supports herb-centered knowledge exploration.

**Fig. 5.**
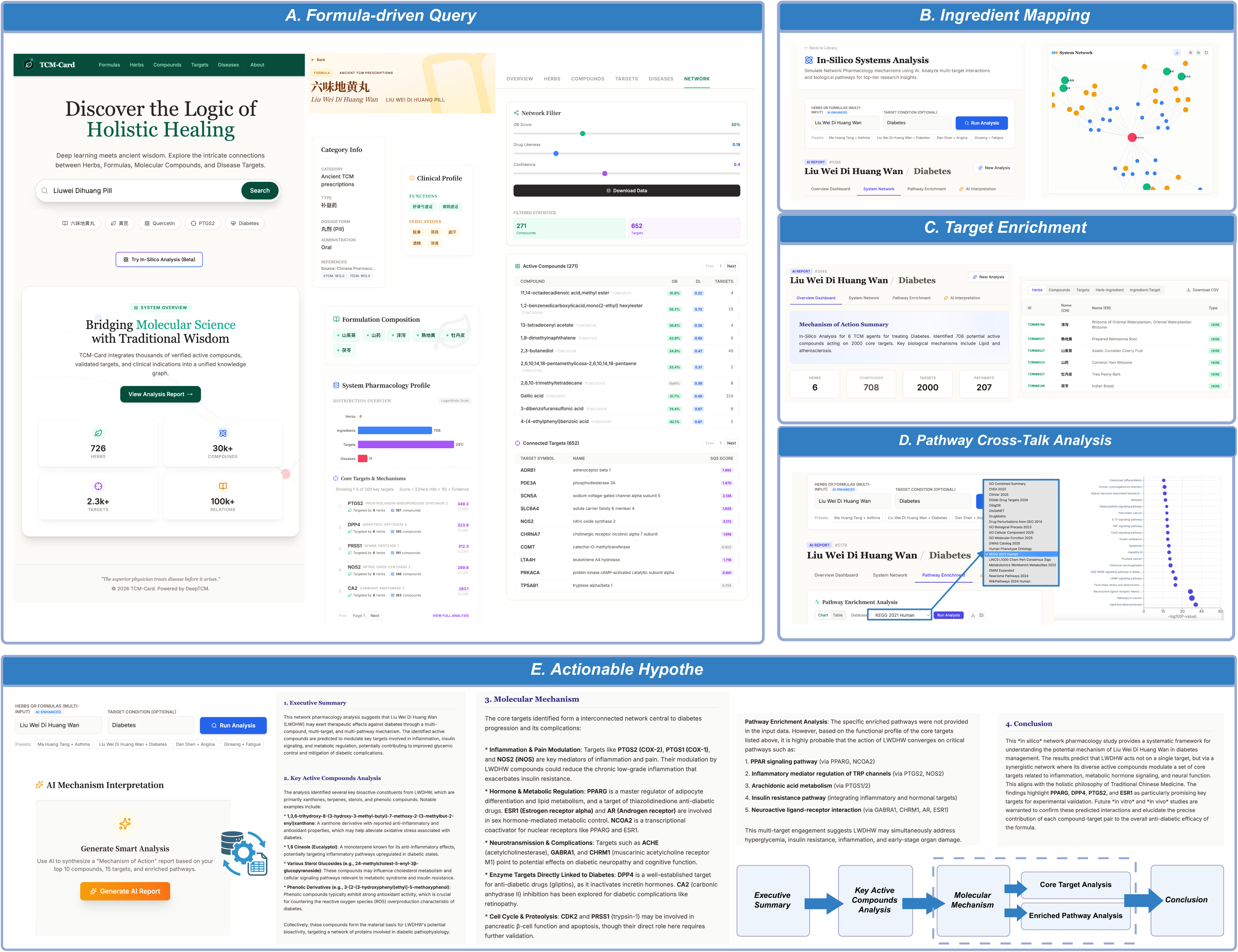
Case-based demonstration of entity-centered exploration in TCMCard using Astragali radix. (A) Overview page for Astragali radix, displaying key information including basic properties, clinical profile, top active ingredients, synergy quality score, and molecular target profile. (B) Entity-specific relationship modules connecting astragali radix to related formulas, compounds, targets, and diseases. (C) Network exploration page with adjustable filters for herb-associated active compounds and connected targets, together with downloadable results.

## Discussion and conclusion

Network pharmacology studies of TCM have long been challenged by heterogeneous, weakly curated interaction data[33]. Although public resources have expanded rapidly, improvements in evidence quality control have lagged behind[19, 21, 34]. As a result, interaction networks often grow in scale without a corresponding increase in biological interpretability. Weakly supported associations can be repeatedly propagated across databases, inflating apparent complexity and obscuring biologically meaningful structure.

Against this background, the main contribution of TCMCard is not simply an integrated resource, but a methodological shift from data aggregation to evidence-structured knowledge organization. Our results show that removing more than 60% of potentially noisy interactions through rigorous curation does not diminish network interpretability. Instead, it improves and clarifies the underlying pharmacological mechanisms. This finding argues against the common assumption in network pharmacology that larger interaction spaces are inherently more informative. In practice, the accumulation of weakly supported edges may inflate topological complexity, adding little analytical value. By contrast, our approach reduces redundancy, suppresses spurious associations, and highlights the core relationships most likely to reflect meaningful pharmacological organization.

This logic also explains why curation must be complemented by confidence modeling. Even after systematic filtering, ingredient-target associations remain inherently highly unequal in evidential strength and therefore should not be treated as analytically equivalent. The MDEI framework addresses this problem by converting disparate evidence types, including bioactivity measurements, literature support, and source provenance, into a unified confidence architecture. Under this framework, each edge is treated as a weighted unit of evidence rather than a simple binary entry. This allows interactions to be ranked, filtered, and interpreted according to their empirical support. Notably, our results reveal a markedly stratified confidence landscape, with only a small fraction of interactions forming a high-confidence core, a prerequisite for robust inference in network pharmacology. This separation is important because downstream tasks such as enrichment analysis, hub prioritization, and predictive modeling are highly sensitive to edge reliability and heterogeneity in confidence.

The impact of this evidence-aware representation becomes especially clear at the topological level. Previous studies have noted that TCM interaction networks often exhibit characteristics similar to those of random graphs due to the presence of numerous noisy edges that blur their internal organization[35]. After pruning low-confidence associations, however, the network topology is markedly reorganized, and TCM-KG exhibits stronger biological network features, including a power-law degree distribution[36], relative robustness to random perturbations, and vulnerability to targeted attacks[37]. These ordered topological features suggest that the TCMCard network is supported by an organized and resilient core architecture. In other words, ingredient-target interactions are not randomly distributed. Instead, they are organized around a structured network backbone. This interpretation aligns with the concept of modular pharmacological regulation in systems pharmacology, which posits that complex therapeutic effects arise from coordinated modulation of multiple functional modules rather than single targets[38]. In this sense, evidence stratification helps reveal system-level organization that may be obscured in noisier interaction spaces and provides a robust foundation for drawing pharmacological conclusions[39].

The observed scaling behavior further suggests that formula complexity is organized rather than unconstrained. This pattern suggests that increasing herb nodes does not produce endless pharmacological novelty, but instead results in more redundancy once a certain point is reached. This means that effective formulas may follow a bounded organizational principle, balancing the range of combinations with manageable overlap. Such a perspective redefines TCM complexity not as arbitrary accumulation, but as a controlled strategy to maintain coordinated multi-target activity while preventing unnecessary network expansion.

The broader value of TCMCard lies in its transformation of an evidence-based interaction landscape into a reproducible analytical environment. To ensure that high-quality data effectively support scientific research, we further developed TCMCard as a cloud-native analysis platform. Unlike traditional databases that primarily offer downloadable tables, TCMCard integrates interactive network visualization and real-time data exploration in a web-based environment, creating a native knowledge ecosystem for TCM research. In conventional database workflows, researchers are often required to download raw datasets and perform extensive preprocessing before conducting network analyses. By contrast, TCMCard enables users to dynamically explore multilayer relationships among formulas, herbs, ingredients, targets, and diseases, monitoring real-time changes in network topology while using MDEI scores to prioritize biologically credible interactions. This interactive framework substantially improves data accessibility and reduces the technical barrier for computational analysis, broadening participation in data-driven research beyond bioinformatics specialists to include clinicians and pharmacologists. Specifically, TCMCard also incorporates an AI-assisted reporting module that summarizes analytical outputs and facilitates result interpretation from network-level results to clearer and more testable biological hypotheses, thereby enhancing the usability of the platform for network pharmacology research. Collectively, the launch of TCMCard marks a shift of TCM databases from static repositories toward interactive research platforms, laying the foundation for a more open, collaborative, and analytically scalable ecosystem in traditional medicine.

Several limitations should still be acknowledged. Although TCMCard significantly enhances data quality through systematic curation and evidence integration, the chemical space covered in the current version remains limited by available public resources. Consequently, low-abundance, unstable, or as-yet-unknown constituents may still be underrepresented. Second, confidence estimation within MDEI is inevitably influenced by the depth, completeness, and bias present in existing bioactivity databases and literature records. Additionally, while this study establishes an evidence-based framework and analytical platform, the predictive usefulness of this resource for AI-driven applications still requires systematic benchmarking and prospective validation[40,41]. Its practical utility will ultimately depend on rigorous evaluation in real-world drug discovery workflows[42]. These limitations do not negate the current findings but highlight the urgent need for more extensive data collection and more scalable evidence integration. Future efforts will focus on broadening both data depth and analytical scope. We plan to go beyond the limits of static public databases by automating the extraction of entity relationships from large-scale biomedical literature and applying deep learning strategies to infer and expand latent chemical and pharmacological space. At the same time, we will continue to enhance the cloud platform through increased integration of multi-omics data, including transcriptomic, proteomic, and single-cell datasets[43], supporting more comprehensive and multidimensional systems-level investigations.

Overall, this study emphasizes the importance of evidence quality control as a key factor in the interpretability of TCM systems pharmacology. By demonstrating how confidence stratification enhances interaction landscapes and clarity, we promote shifting from passive database growth to evidence-aware knowledge structures. TCMCard serves not only as a benchmark resource but also as a foundational method for identifying which pathways within complex networks are sufficiently supported to allow for rigorous biological interpretation.

## Supporting information

Hierarchical quality distribution and semantic value of relational tiers

## Acknowledgement

The authors thank the Beijing Natural Science Foundation and the National Natural Science Foundation of China for their support.

## Author Contributions

**Yuqi Wang**: Writing – original draft, Formal analysis, Conceptualization, Methodology. **Wenqing Dong**: Visualization, Writing – review & editing, Investigation. **Junkai Yao**: Visualization, Writing – review & editing, Investigation. **Kai Wang**: Investigation, Formal analysis, Writing – review & editing. **Leyuan Zhang**: Data curation, Investigation, Writing – review & editing. **YuanRong Wang**: Writing – review & editing, Visualization, Investigation. **ShanShan Guo**: Writing – review & editing, Investigation. **Haorui Li**: Writing – review & editing, Methodology. **Hengjia Cai**: Writing – review & editing, Investigation. **Xiting Wang**: Writing – review & editing, Conceptualization, Methodology, Funding acquisition. **Yu Li**: Writing – review & editing, Supervision, Project administration, Funding acquisition, Conceptualization.

## Supplementary materials

Table S1. Comparison of representative TCM resources and TCMCard

Table S2. Ingredient-Target Relationship Details (MDEI Model)

Table S3. Entity Quality Distribution

Table S4. Confidence distribution across relationship types

## Statement of Ethics

A Statement of Ethics is not applicable because this study used only publicly available literature and database resources and did not involve human participants or animals.

## Data Availability

The data supporting this study are available through the TCMCard website at https://www.tcmcard.com.

