## Supplementary material for "TCMCard: A High-Confidence Digital Infrastructure for Traditional Chinese Medicine Quantified by Multi-Dimensional Evidence Integration": Hierarchical quality distribution and semantic value of relational tiers

Table 1. Hierarchical quality distribution and semantic value of relational tiers

| **Relation type** | **Total edges** | **Mean score** | **Descriptive values** |
| --- | --- | --- | --- |
| **H-I (Herb-Ingredient)** | 56678 | 0.750 | Reflects high-confidence relationships between herbs and their chemical constituents. |
| **F-H (Formula-Herb)** | 16303 | 0.700 | Captures the classical formula-herb composition relationships in traditional TCM prescriptions. |
| **H-D (Herb-Disease)** | 10292 | 0.600 | Represents clinically supported associations between medicinal materials and specific diseases. |
| **F-D (Formula-Disease)** | 9464 | 0.600 | Indicates clinical associations between classical formulas and specific diseases. |
| **T-D (Target-Disease)** | 15950 | 0.600 | Maps disease-associated molecular targets and therapeutic markers. |
| **I-D (Ingredient-Disease)** | 98005 | 0.500 | Suggests potential pharmacological relevance of individual ingredients to disease phenotypes. |
| **I-T (Ingredient-Target)** | 150183 | 0.460 | Encapsulates the complex interactome between chemical compounds and molecular targets. |
