## Supplementary materials for "TCMCard: A High-Confidence Digital Infrastructure for Traditional Chinese Medicine Quantified by Multi-Dimensional Evidence Integration"

**Table S1. Comparison of representative TCM resources and TCMCard**

| **Resources** | **Year** | **Core design goal** | **Primary knowledge scope** | **Construction strategy** | **Distinctive features** | **Dimension not emphasized relative to TCMCard** |
| --- | --- | --- | --- | --- | --- | --- |
| TCM-ID | 2006 | Build an early comprehensive information resource for TCM prescriptions, herbs, and ingredients | prescriptions, constituent herbs,  herbal ingredients, molecular structure and functional properties of active ingredients, therapeutic and side effects, clinical indication | Manually collected information on TCM in books and other printed sources | Early efforts toward exploring TCM for new theories in drug discovery. | A foundational descriptive resource without information about the connections between ingredients and targets |
| TCM Database@Taiwan | 2011 | Enable virtual screening or molecular simulation of a 3D small molecular structure database of TCM | TCM ingredients / pure compounds / 2D–3D molecular structures | Manual collection from Chinese medical texts, dictionaries, and scientific publications | Large downloadable 2D/3D TCM compound library | An early structure-oriented TCM resource focused on compound collection and virtual screening, with limited support for formula-centric curation, evidence integration, and mechanistic network interpretation |
| TCMID | 2013 / 2018 (v2.0) | Bridge TCM and modern biomedicine by integrating multi-layer TCM entities into modern Western medicine | Prescriptions, herbs, ingredients, targets, drugs, diseases, and connections between them; TCMID 2.0 further extends to herbal MS spectra, ingredient MS spectra, and PPIs. | Manually collected through literature mining and integration of heterogeneous resources | Early large-scale integrative framework linking prescriptions, herbs, ingredients, targets, and diseases | Primarily a breadth-oriented integrative resource rather than a high-fidelity curated benchmark |
| TCMSP | 2014 | Build an efficient systems pharmacology platform, integrating pharmacochemistry, ADME properties, drug-likeness, drug targets, associated diseases, and interaction networks to accelerate the drug discovery | Herbs, ingredients, targets, diseases, ADME-related features, and drug targets and diseases of each active compound | Integrate large-scale structures data with manually curated information in the Chinese pharmacopoeia; incorporating diverse sources for active compound screening | Strong ADME-driven screening and establishment of the compound-target, target-disease networks | Primarily optimized for ADME-guided screening and hypothesis generation, rather than evidence stratification |
| BATMAN-TCM | 2016 / 2024 (v2.0) | Build an integrative bioinformatics resource for known and predicted TTIs to support mechanism study, and in v2.0, further enable omics-guided discovery of candidate TCM ingredients/herbs for complex diseases | Ingredients, target proteins, herbs, formulas, and ingredient-target interactions  Known and predicted TTIs linking TCM ingredients to target proteins, together with herbs, formulas, pathways/functions, and disease-related annotations; v2.0 substantially expands coverage | Manual curation of known TTIs from prior BATMAN-TCM, other TCM databases including DrugBank, KEGG, TTD, HIT, and HERB, and large-scale prediction of high-confidence TTIs | Strong proteome-scale TTI coverage, joint presentation of known and predicted interactions, traceable evidence pages for known and predicted TTIs, and practical browse/download/API support. | Prediction-oriented and optimized for interaction discovery rather than evidence-prioritized denoising and confidence-stratified mechanistic interpretation |
| TCM-Mesh | 2017 | Integrate a database and a data-mining system for one-stop network pharmacology analysis of TCM preparations | Herbs, compounds, genes, diseases, gene-disease associations, pairs of gene interactions, side effects, toxicity | Large-scale integration and filtration of heterogeneous databases (including TCMID, STITCH, STRING, OMIM, GAD, TOXNET, and SIDER), identifier unification through canonical SMILES and gene mapping, redundancy trimming, and score-threshold-based network construction | Broad network pharmacology coverage with unusually broad inclusion of safety information; supports herb/compound/gene/disease queries, downloadable reports, and visualization of compound-gene-disease chains | Filtering in TCM-Mesh is primarily redundancy/format-oriented rather than clinically informed and evidence-prioritized, and its scoring framework is relatively primitive, limiting high-reliability mechanistic interpretation |
| YaTCM | 2018 | Provide a comprehensive TCM database equipped with an integrated TCM toolkit for drug discovery and mechanism exploration | Prescriptions, herbs, ingredients, targets, pathways, predicted targets | Manual integration of heterogeneous resources (including TCMID, Database@Taiwan, TCMSP, TTD, ChEMBL, KEGG, books, and PubMed text mining), duplicate removal, KEGG/ChEMBL mapping, ADMET descriptor calculation, and regular updating through repeated text mining and manual curation | An analysis toolkit for similarity/substructure searches, MV-SEA for predicting protein targets, network analysis, pathway analysis, and identification of functionally similar herb pairs | Tool-rich and analysis-oriented, but not intended as a high-fidelity benchmark for confidence-stratified formula interpretation |
| ETCM | 2019 / 2023 (v2.0) | Build a comprehensive encyclopaedia of TCM (with the habitat and quality control information of herbs, and drug-likeness information of ingredients) | Formula, Chinese patent drugs, Chinese medicinal materials, ingredients, confirmed or potential drug targets, and related diseases. | Manual integration and standardization of pharmacopoeial/clinical sources with public databases (including ChEMBL, PubChem, DrugBank, KEGG, HPO, OMIM, DisGeNET, ORPHANET, Reactome, HPRD, MINT, IntAct and DIP); ETCM v2.0 further incorporated BindingDB-supported target identification and expanded curated formula/drug associations | Standardization and annotation richness, with pharmacopoeia-based quantitative information of marker ingredients, habitat, and TCM-property data, formula/patent-drug coverage, similarity evaluation, and enhanced network visualization | Strong in standardization, quality-control annotation, and clinical/formula coverage, its mechanistic association layer still depends mainly on similarity- and enrichment-based linkage |
| SymMap | 2019 | Integrate TCM with modern medicine through symptom mapping, linking phenotypes with herbs and molecular mechanisms | TCM symptoms, modern symptoms, herbs, ingredients, diseases, targets | Manual curation and standardization of TCM symptoms from the Chinese Pharmacopoeia, expert mapping to UMLS/ MeSH/ SIDER symptoms, and integration of TCMID, TCMSP, TCM-ID, HIT, HPO, DrugBank, NCBI Gene, OMIM, and Orphanet with statistically inferred indirect associations | Expert-curated TCM symptom-modern medicine symptom mapping and a phenotype-enhanced heterogeneous network | Phenotype-oriented, symptom-centered, and statistically inferred, but less focused on clinically grounded formula curation |
| HERB | 2021 / 2025 (v2.0) | Build an experiment- and evidence-guided TCM database, later expanded to integrate clinical and experimental evidence for mechanism study and drug discovery | Herbs, ingredients, targets, diseases, high-throughput experiments, curated references, and CMap-linked modern drugs; HERB 2.0 further integrates meta-analyses and clinical trials | Integrated and de-redundant data from SymMap, TCMID 2.0, TCMSP, TCM-ID, and the National Database for Chemical Composition in TCM; re-analyzed GEO pharmacotranscriptomics data and mapped them to CMap; manually curated PubMed evidence, and in HERB 2.0 further incorporated formulae from Pharmacopoeia/CFDA/ancient classics, clinical evidence from ClinicalTrials.gov and PROSPERO | Strong transcriptomic and literature/clinical-backed evidence integration, and knowledge-graph/network-based exploration in one platform | Emphasizes pharmacotranscriptomic, literature, and clinical evidence integration, but still retains heterogeneity in relationship confidence and formula-level standardization |
| SuperTCM | 2021 | Build a biocultural database combining biochemical genetic pathways and historical linguistic data of Chinese Materia Medica for drug development. | Herbs, accepted botanical names, botanical synonyms, common plant names, recipes, ingredients, targets, diseases, pathways, and KEGG global maps | Anchored on the Chinese Pharmacopoeia, MPNS/NCBI botanical normalization, extracted data from multiple sources (including Chinese Herbal Medicine: Formulas & Strategies, Merged Herb, TCMID 2.0, TCM Database@Taiwan, CMAUP, TCM-Mesh, TM-MC, and ETCM, and retained high-confidence human ingredient–target pairs from ChEMBL for KEGG/TTD/ICD-10-CM linkage. | Botanical-source standardization, ICD-10 disease linkage, and pathway visualization. | Its quality-control logic strategy is centered primarily on botanical normalization rather than formula-centered standardization across all entity layers. |
| HIT 2.0 | 2011 / 2022 (v2.0) | Curate dataset focusing on Herbal Ingredients’ targets covering PubMed literature 2000–2020. | Literature-described targets, herbal ingredients, and herbal ingredient-target activity pairs | Literature-curated from PubMed abstracts: HIT 1.0 covered 2000–2010 and HIT 2.0 extended this to 2000–2020 with TCM-ID/CAS-based ingredient normalization, NLP-assisted screening, and multi-curator manual verification. | A focused literature-curated resource for herbal ingredient-target relationships, with explicit interaction-type annotation, literature-based quality indicators, and, in HIT 2.0, update-oriented target mining and user-assisted online curation. | Although literature curated, the interaction evidence remains heterogeneous in assay type, validation depth, and network-level confidence stratification |
| TCMBank | 2023 | Build the largest systematic free TCM database and support AI-assisted drug discovery | Herbs, ingredients, targets, diseases, pairwise relationships, 3D ingredient structures, and external cross-references | Integrated books, literature, and major TCM databases (including TCMID, TCMSP, SymMap, TCM-ID, HERB, ETCM), then standardized target/disease records with public biomedical resources, with AI-assisted document mining and double manual verification for continuous updating | Large-scale integrated resource with continuous literature updating, 3D structures, and AI-oriented exploration. | Broad and continuously updated, its quality control is mainly entity- and linkage-level, rather than formula-centered standardization, evidence-prioritized curation, and confidence-stratified refinement as in TCMCard |
| **TCMCard** | 2026 (this study) | Build a formula-centered, evidence-weighted, confidence-stratified TCM knowledge graph and web platform for standardized network pharmacology, knowledge retrieval, and predictive inference | Formulas, herbs, ingredients, targets, diseases, and confidence-stratified interaction relationships | Formula-centered curation from Chinese Pharmacopoeia and classical prescriptions, ingredient refinement by pharmacopoeial cross-referencing and cheminformatic QC, interaction validation/expansion using ChEMBL and PubChem, disease integration, and MDEI-based confidence stratification | Confidence-aware denoising with removal of over 60% low-confidence links, a high-confidence interaction core, and integrated retrieval, network analysis, predictive inference, and AI-assisted reporting | Not applicable |

TCM: traditional Chinese medicine; MS: mass spectrometry; PPI/PPIs: protein–protein interaction(s); TTI/TTIs: TCM ingredient–target protein interaction(s); OMIM: Online Mendelian Inheritance in Man; ADME: absorption, distribution, metabolism, excretion; NLP: natural language processing; UMLS: Unified Medical Language System; SIDER: Side Effect Resource; CMap: Connectivity Map; CFDA: China Food and Drug Administration; PROSPERO: International Prospective Register of Systematic Reviews; GEO: Gene Expression Omnibus; MPNS: Medicinal Plant Names Services; API: application programming interface; ICD-10-CM: International Classification of Diseases, Tenth Revision, Clinical Modification; CAS: Chemical Abstracts Service; MDEI: multi-dimensional evidence integration

Table S2. Ingredient-Target Relationship Details (MDEI Model)

| **Evidence Type** | **Count** | **Percentage** |
| --- | --- | --- |
| Experimental activity | 15,874 | 10.6% |
| Literature support | 15,529 | 10.3% |
| Database only | 111,599 | 74.3% |
| External | 7,181 | 4.8% |

Table S3. Entity Quality Distribution

| **Entity type** | **Total** | **High(≥0.8)** | **Medium(0.50-0.80)** | **Low(<0.5)** | **Mean score** |
| --- | --- | --- | --- | --- | --- |
| **Formula** | 1,774 | 1,760 (99.2%) | 14 (0.8%) | 0 | 0.882 |
| **Herb** | 748 | 730 (97.6%) | 18 (2.4%) | 0 | 0.928 |
| **Ingredient** | 30,787 | 30,787 (100%) | 0 (0%) | 0 | 0.926 |
| **Target** | 10,704 | 10,684 (99.8%) | 20 (0.2%) | 0 | 0.978 |
| **Disease** | 2,459 | 2,412 (98.1%) | 47 (1.9%) | 0 | 0.857 |
| **Total** | 46,472 | 46,373 (99.8%) | 99 (0.2%) | 0 | 0.914 |

Table S4. Confidence distribution across relationship types

| **Relationship Type** | **Total** | **Highest (≥0.90)** | **High (0.75-0.89)** | **Medium (0.55-0.74)** | **Low (<0.55)** | **Mean Score** |
| --- | --- | --- | --- | --- | --- | --- |
| **Formula-Herb** | 16,303 | 0 (0.0%) | 0 (0.0%) | 16,303 (100.0%) | 0 (0.0%) | 0.700 |
| **Herb-Ingredient** | 56,678 | 0 (0.0%) | 0 (0.0%) | 56,678 (100.0%) | 0 (0.0%) | 0.750 |
| **Ingredient-Target** | 150,183 | 6,818 (4.5%) | 16,237 (10.8%) | 15,529 (10.3%) | 111,599 (74.3%) | 0.460 |
| **Herb-Disease** | 10,292 | 0 (0.0%) | 0 (0.0%) | 0 (0.0%) | 10,292 (100.0%) | 0.600 |
| **Formula-Disease** | 9,464 | 0 (0.0%) | 0 (0.0%) | 0 (0.0%) | 9,464 (100.0%) | 0.600 |
| **Ingredient-Disease** | 98,005 | 0 (0.0%) | 0 (0.0%) | 0 (0.0%) | 98,005 (100.0%) | 0.500 |
| **Target-Disease** | 15,950 | 0 (0.0%) | 0 (0.0%) | 0 (0.0%) | 15,950 (100.0%) | 0.600 |
| **Total** | 356,875 | 6,818 (1.9%) | 16,237 (4.5%) | 88,510 (24.8%) | 245,310 (68.7%) | 0.601 |
